# Cerebral Atrophy in Amyotrophic Lateral Sclerosis Parallels the Pathological Distribution of TDP43

**DOI:** 10.1101/2020.02.18.954883

**Authors:** Mahsa Dadar, Ana Laura Manera, Lorne Zinman, Lawrence Korngut, Angela Genge, Simon J. Graham, Richard Frayne, D. Louis Collins, Sanjay Kalra

**Affiliations:** McConnell Brain Imaging Centre, Montreal Neurological Institute, McGill University, Montreal, Quebec, Canada; Sunnybrook Health Sciences Centre, University of Toronto, Toronto, Ontario, Canada; Departments of Radiology and Clinical Neurosciences, Hotchkiss Brain Institute, University of Calgary, and Seaman Family MR Research Centre, Foothills Medical Centre, Calgary, Alberta, Canada; Montreal Neurological Institute and Hospital, McGill University, Montreal, Quebec, Canada; Neuroscience and Mental Health Institute; Department of Medicine, Division of Neurology; University of Alberta, Edmonton, Canada

**Keywords:** Amyotrophic lateral sclerosis, deformation-based morphometry, magnetic resonance imaging, mixed effects modeling

## Abstract

Amyotrophic lateral sclerosis (ALS) is a neurodegenerative disease characterized by a preferential involvement of both upper and lower motor neurons. Evidence from neuroimaging and post-mortem studies confirms additional involvement of brain regions extending beyond the motor cortex. The aim of this study was to assess the extent of cerebral disease in ALS cross-sectionally and longitudinally, and to compare the findings with a recently proposed disease-staging model of ALS pathology. Deformation-based morphometry (DBM) was used to identify the patterns of brain atrophy associated with ALS and to assess their relationship with clinical symptoms. Longitudinal T1-weighted MRI data and clinical measures were acquired at baseline, 4 months, and 8 months, from 66 ALS patients and 43 age-matched controls who participated in the Canadian ALS Neuroimaging Consortium (CALSNIC) study. Whole brain voxel-wise mixed-effects modelling analysis showed extensive atrophy patterns differentiating ALS patients from the normal controls. Cerebral atrophy was present in the motor cortex and corticospinal tract, involving both GM and WM, and to a lesser extent in non-motor regions. More specifically, the results showed significant bilateral atrophy in the motor cortex, the corticospinal tract including the internal capsule and brainstem, with an overall pattern of ventricular enlargement; along with significant progressive longitudinal atrophy in the precentral gyrus, frontal and parietal white matter, accompanied by ventricular and sulcal enlargement. Atrophy in the precentral gyrus was significantly associated with greater disability as quantified with the ALS Functional Rating Scale-Revised (ALSFRS-R) (p<0.0001). The pattern of atrophy observed using DBM was consistent with the Brettschneider’s four stage pathological model of the disease. Deformation based morphometry provides a sensitive indicator of atrophy in ALS, and has potential as a biomarker of disease burden, in both gray and white matter.

## Introduction

Amyotrophic lateral sclerosis (ALS) is a progressive neurodegenerative disorder, with a highly selective involvement of the upper motor neurons (UMN) in the cortex and lower motor neurons (LMN) in the brainstem and spinal cord (Brownell *et al.*, 1970; Martin and Swash, 1995). This characteristic neuronal atrophy in the motor cortex is also accompanied by degeneration in the white matter tracts, most severely in the precentral areas and to a lesser extent elsewhere (Kushner *et al.*, 1991). Beyond the motor pathways, degeneration in the frontal and temporal lobes is the substrate for the cognitive deficits present in ALS (Strong, 2001).

Onset of ALS symptoms is variable, and can include involvement of limb, bulbar or respiratory muscles or cognitive impairment. Spasticity, hyperflexia, and weakness result from UMN degeneration (Gordon, 2013), whereas fasciculations, cramps and muscle wasting result from LMN degeneration (Hardiman *et al.*, 2017). Cognitive dysfunction also affects more than half of the patients, with evident frontotemporal dementia (FTD) in 15% (Wilson *et al.*, 2001; Ringholz *et al.*, 2005; Landau *et al.*, 2012). Approximately 30% of all patients have evidence of executive dysfunction at onset, and FTD is one of the presenting features in 13% of incident cases (Hardiman *et al.*, 2017).

Previous structural MR imaging studies have demonstrated brain changes arising from ALS, although with relatively poor sensitivity and specificity (Kato *et al.*, 1993; Oba *et al.*, 1993; Waragai, 1997; Andreadou *et al.*, 1998; Hecht *et al.*, 2001; Charil *et al.*, 2009). Voxel Based Morphometry (VBM) studies report a trend of grey matter loss in the motor cortex, and in frontotemporal regions especially for patients with cognitive deficits (Chang *et al.*, 2005; Agosta *et al.*, 2007, 2009). These regional studies have generally been performed in relatively small samples and have not yet produced conclusive evidence of motor cortex or corticospinal tract atrophy. To date, no MRI study has been able to provide a comprehensive picture of the brain regions that are involved in ALS, and conventional MRI still remains a tool to rule out disorders that mimic ALS rather than to confirm diagnosis or quantify disease burden.

In this study, we have used deformation-based morphometry (DBM) to quantify the patterns of disease-related brain changes in ALS. Unlike voxel-based morphometry (VBM), DBM does not rely on a prior tissue classification step, which might introduce errors into the measurements, particularly when the population under study presents with white matter hyperintensities on T2-weighted images. This a relevant concern in ALS studies, where there is evidence of white matter tract alterations appearing as T2-weighted hyperintensities in a proportion of patients (Hecht *et al.*, 2001). Such errors might create systematic biases in VBM measurements. In addition, the image processing tools used in this study have been designed for use in multi-center datasets across different MRI systems, and are able to accommodate between-site variabilities. Taking advantage of these superior methodological tools, the primary goal was to quantify accurately the patterns of disease-related brain changes in ALS and investigate the associations between such changes and clinical symptoms.

In addition, these *in vivo* changes were investigated in relation to the pathological staging system of Brettschneider *et al.* (Brettschneider *et al.*, 2013), that characterizes ALS in four stages based on the regional accumulation of transactive response DNA-binding protein 43kDa (TDP-43). Stage one is characterized by involvement of the primary motor cortex, alpha motor neurons in the ventral horn of the spinal cord, and brainstem motor nuclei of cranial nerves V, VII, and X-XII. Stage two adds involvement of the prefrontal neocortex, reticular formation and the inferior olivary complex. In stage three, TDP-43 deposition is now present more extensively in the prefrontal and postcentral neocortex and striatum. Stage four additionally involves the anteromedial portions of the temporal lobe, including the hippocampal formation. It was investigated whether brain atrophy, estimated by DBM analysis, followed the Brettschneider stages of ALS pathology.

## Materials and Methods

### Participants

Data used in this study included longitudinal (baseline, month 4, and month 8) MRI and clinical measurements of 66 ALS patients (N_Baseline_ = 64, N_Month4_ = 44, N_Month8_ = 24) as well as 43 age-matched healthy controls (N_Baseline_ = 42, N_Month4_ = 32, N_Month8_ = 21) from the Canadian ALS Neuroimaging Consortium (CALSNIC) (http://calsnic.org/, ClinicalTrials.gov NCT02405182). Each participating centre in CALSNIC followed identical standard operating procedures for clinical evaluations and harmonized acquisition protocols for brain imaging. Data from four sites were included: University of Alberta, University of Calgary, University of Toronto, and McGill University. All participants gave written informed consent and the study was approved by the health research ethics boards at each of the participating sites. Participants were included if they had sporadic or familial ALS, and met El Escorial criteria for possible, probable, probable laboratory supported, or definite ALS (Brooks *et al.*, 2000). Participants were excluded if they had a history of other neurological or psychiatric disorders, prior brain injury, or respiratory impairment resulting in an inability to tolerate the MRI protocol.

Clinical evaluations included global measures of disease status, upper motor neuron dysfunction, and cognitive batteries. Disability was assessed with the ALS Functional Rating Scale-Revised (ALSFRS-R), and the annualized disease progression rate was estimated using the formula: (48-ALSFRS-R)/symptom duration. Finger and foot tapping rates (left and right) were measured and averaged as indicators of UMN function. Cognitive function was assessed using tests for verbal fluency (letter F), digit span, and the Edinburgh Cognitive and Behavioral ALS Screen (ECAS) scores (Abrahams *et al.*, 2014).

All imaging data were acquired on 3 Tesla MRI systems. University of Alberta and McGill University acquired data using Prisma and TimTrio Siemens systems, respectively, and University of Toronto and University of Calgary acquired data using General Electric Healthcare (Discovery MR750) systems. The MRI acquisition protocol included three-dimensional (3D) T1-weighted scans acquired at 1 mm^3^ isotropic resolution. Acquisition parameters were identical for the Siemens systems, which used a Magnetization Prepared RApid Gradient Echo (MPRAGE) sequence with repetition time [TR] = 2300 ms, echo time [TE] = 3.43 ms, inversion time [TI] = 900 ms, flip angle = 9°, field of view [FOV] = 256 mm × 256 mm. The two General Electric systems used identical parameters as well for an inversion recovery-prepared fast spoiled gradient-recalled echo (FSPGR) imaging sequence with TR = 7.4 ms, TE = 3.1 ms, TI = 400 ms, flip angle = 11°, FOV = 256 mm × 256 mm.

### MRI Processing

All T1-weighted MRI data were pre-processed using the Medical Imaging Network Common dataform (MINC) toolkit of the Montreal Neurological Institute (MNI), publicly available at https://github.com/BIC-MNI/minc-tools, using the following steps: 1) denoising (Coupe *et al.*, 2008); 2) intensity inhomogeneity correction (Sled *et al.*, 1998); and 3) image intensity normalization into intensity range [0-100] according to a linear histogram matching algorithm. All images were first linearly (Dadar *et al.*, 2018*a*) and then nonlinearly (Avants *et al.*, 2008) registered to an average template recognized by the International Consortium for Brain Mapping (ICBM), MNI-ICBM152 (Fonov *et al.*, 2009). The quality of the registrations was visually assessed, confirming that all imaging data were accurately registered to the template.

Using MINC tools, DBM maps were obtained by computing the Jacobian of the estimated non-linear deformation field. DBM maps reflect the local differences between the average MNI-ICBM152 template (Fonov *et al.*, 2009) and images of a given participant, with values larger than one indicating expansion (e.g. larger sulci or ventricles) and values smaller than one indicating shrinkage (e.g. atrophy). Voxel-wise DBM maps were used to assess the global brain differences between controls and ALS patients. Mean DBM values across regions of interest were used to assess the relationship between regional atrophy and clinical measurements.

### Statistical Analyses

To investigate brain changes related to ALS in relation to the confounds of normative aging as well as sex differences, a regression procedure was developed involving the images from the age-matched controls, as similarly performed in previous studies (La Joie *et al.*, 2012; Moradi *et al.*, 2015; Zeighami *et al.*, 2017; Brown *et al.*, 2019)). The following voxel-wise mixed effects models were estimated based on the controls:

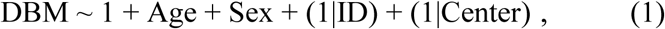

where DBM indicates the DBM value at a certain voxel in the brain; Age indicates the participant’s age at the time of imaging; Sex (Male/Female) is a categorical variable; and ID and Center are categorical random variables, indicating the control participant’s ID and the data acquisition center. Based on the model estimates, normative aging and sex effects were then regressed out from the patients. The following mixed effects models were used to estimate the ALS-specific brain changes in the patients, accounting for confounds:

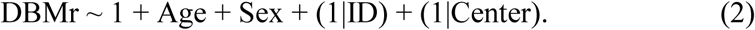

Here, DBMr are the residualized DBM values (after regressing out age and sex based on controls, i.e. eq. 1), also referred to in the literature as a W-score map (La Joie *et al.*, 2012; Brown *et al.*, 2019). The resulting maps were corrected for multiple comparisons using False Discovery Rate (FDR) with a significance threshold of 0.05.

In addition, mean DBM values were calculated for the corticospinal tract mask obtained from Yeh et al. (Yeh *et al.*, 2018), as well as the frontal lobe, frontotemporal lobe, precentral gyrus, and dorsolateral prefrontal cortex masks obtained from a grey matter atlas for the MNI-ICBM152 template (Manera *et al.*, 2019). The average values were used in the following region-of-interest (ROI) analyses:

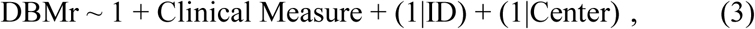

where Clinical Measure indicates either categorical dichotomized scores (e.g. Median Progression Rate [Fast/Slow]) or continuous values (e.g. ALSFRS-R [0-48]). Table 1 indicates the list of the regions and measures of interest tested. Results were corrected for multiple comparisons using FDR with a significance threshold of 0.05.

**Table 1.** Regions and clinical measures of interest assessed in this study.

| Measure | UMN Burden | Progression Rate | Cognitive status | Executive Impairment | Site of Onset | Average Tapping Rate | ALSFRS-R |
| --- | --- | --- | --- | --- | --- | --- | --- |
| Corticospinal Tract | X | X |  |  | X | X | X |
| Precentral Gyrus | X | X |  |  | X | X | X |
| Frontotemporal Lobe |  |  | X |  |  |  | X |
| Dorsolateral Prefrontal Cortex |  |  |  | X |  |  |  |

## Results

Table 2 summarizes the demographic information as well as the clinical scores for the participants used in this study.

**Table 2.** Descriptive baseline statistics for the CALSNIC subjects enrolled in this study. Data are number of participants in each category (N), their percentage of the total population in each category (%), and mean ± standard deviation (SD) of key demographic variables. CALSNIC= Canadian ALS Neuroimaging Consortium. UMN= Upper Motor Neuron. ECAS= Edinburgh Cognitive and Behavioral ALS Screen. ALFRS-R= Amyotrophic Lateral Sclerosis Functional Rating Scale-Revised. Significant differences (p<0.05) are displayed in bold font. Dichotomization details: ECAS: Total score, using 2 SD based on controls as cut-off. ALSexi: Total score, using 2 SD based on controls as cut-off. UMN Burden: above/below (“High/Low”) median value. Progression Rate: above/below (“High/Low”) median value.

|  | Control | ALS |
| --- | --- | --- |
| Participants (N) | 42 | 66 |
| Female (N) | 21 (51%) | 24 (36%) |
| Age (years) | 55.03 ± 8.52 | 57.98 ± 10.84 |
| Education (years) | 16.12 ± 2.61 | 14.80 ± 3.26 |
| Symptom Duration (years) | - | 3.32 ± 3.03 |
| Site of Onset (Bulbar/Limb) | - | 11/52 (17/78 %) |
| UMN Burden (High/Low) | - | 20/38 (30/58 %) |
| Progression Rate (High/Low) | - | 35/29 (53/44%) |
| ECAS: Cognitive Status (Impaired/Normal) | - | 30/33 (46/50%) |
| ALSexi: Executive Impairment (Impaired/Normal) | - | 26/37 (40/56%) |
| Average Tapping Rate | **46.13 ± 12.37** | **30.19 ± 15.20** |
| ALSFRS-R | - | 38.3 ± 5.2 |

Figure 1 shows the t-statistics maps reflecting the significant brain patterns (after FDR correction) in the ALS cohort after regressing out the normal expected Age and Sex related changes based on the control group (i.e. the intercept term in model 2). Warm colors indicate areas that are significantly enlarged in the patients in comparison with the controls (e.g. in the sulci and ventricular regions) and cold colors indicate areas of additional shrinkage (i.e. atrophy) in the patients. The results show significant bilateral atrophy in the motor cortex, the corticospinal tract including the internal capsule and brainstem, along with an overall pattern of ventricular enlargement. In addition, significant white matter atrophy was found in anterior cingulate and bilateral posterior parietal areas.

**Fig. 1.**
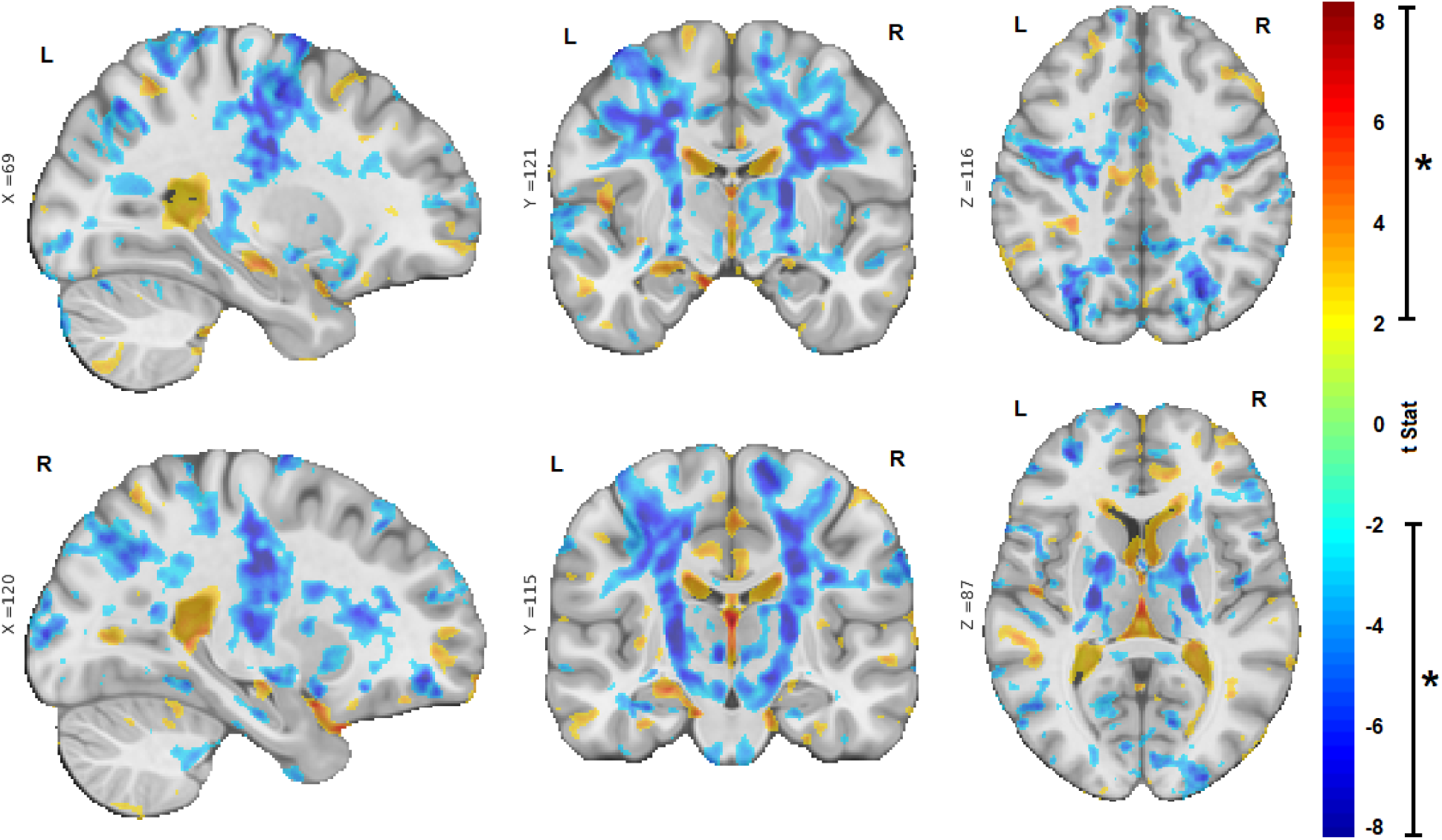
Sagittal, coronal, and axial slices showing the t-statistic maps reflecting the significant patterns of brain volume changes in the ALS cohort (corrected for expected age and sex changes using eq. 1). Warm colors indicate enlargement (e.g. ventricular and sulcal regions) and cold colors indicate shrinkage of the tissue (i.e. atrophy). X, Y, and Z values indicate MNI coordinates for the displayed slice.

Similarly, Figure 2 shows the t-statistics maps reflecting the significant brain changes with age (after FDR correction) only in the ALS cohort. As normative aging has already been regressed out from the model, these remaining changes are the ongoing additional changes specific to the disease occurring during imaging conducted longitudinally over time. The results show additional atrophy in the precentral gyrus that is more pronounced on the right, as well as diffuse atrophy in frontal and parietal white matter and ventricular and sulcal enlargement.

**Fig. 2.**
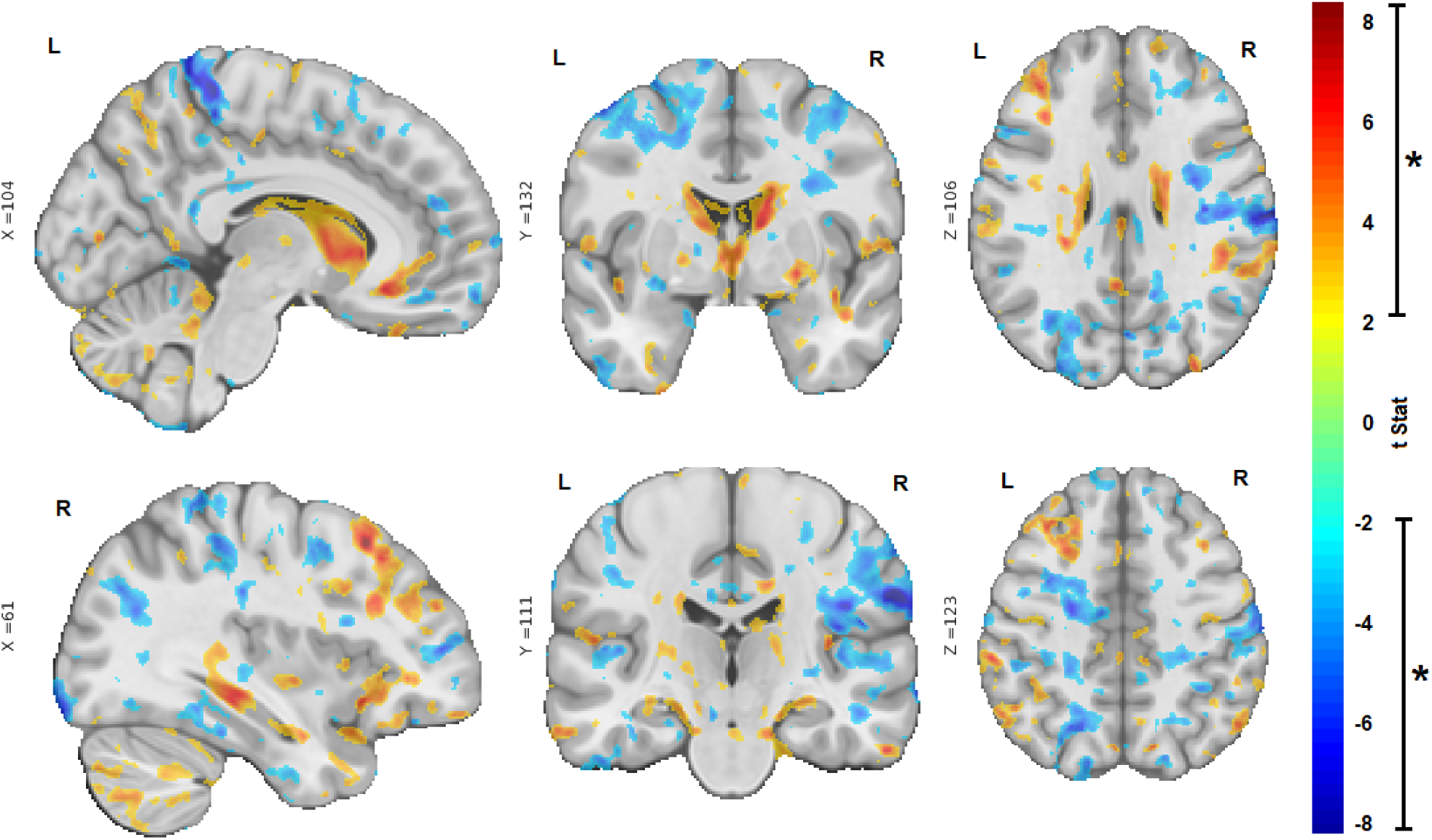
Sagittal, coronal, and axial slices showing the t-statistic maps reflecting the significant brain volume changes with age (i.e. over time) in the ALS cohort. Warm colors indicate enlargement (e.g. ventricular and sulci regions) and cold colors indicate shrinkage of the tissue (i.e. atrophy). X, Y, and Z values indicate MNI coordinates for the displayed slice.

Table 3 shows the associations between clinical measures and regional atrophy. Mean DBM values in the precentral gyrus were significantly and positively associated with ALSFRS-R, indicating that a lower DBM value (i.e. greater atrophy) related to a lower score in ALSFRS-R (i.e. greater disability). Similarly, mean DBM value in the corticospinal tract was positively associated with ALSFRS-R, but this relationship did not survive multiple comparison correction (uncorrected p-value = 0.03). There was also a trend towards an association between lower mean DBM value in the dorsolateral prefrontal cortex and greater executive impairment (uncorrected p-value = 0.08).

**Table 3.** Summary of the results of the mixed effects models of associations between regional DBM values and clinical measures. DBM= Deformation Based Morphometry. CALSNIC= Canadian ALS Neuroimaging Consortium. UMN= Upper Motor Neuron. ECAS= Edinburgh Cognitive and Behavioral ALS Screen. ALFRS-R= Amyotrophic Lateral Sclerosis Functional Rating Scale-Revised. Significant results after correction for multiple comparisons are displayed in bold font.

| Region | t-stat | p-value |
| --- | --- | --- |
| UMN Burden |  |  |
| Corticospinal Tract | -0.03 | 0.97 |
| Precentral Gyrus | -0.33 | 0.74 |
| Progression Rate |  |  |
| Corticospinal Tract | 1.18 | 0.24 |
| Precentral Gyrus | 1.29 | 0.20 |
| Cognitive Status |  |  |
| Frontotemporal Lobe | 1.39 | 0.17 |
| Executive Impairment |  |  |
| Dorsolateral Prefrontal Cortex | -1.72 | 0.08 |
| Site of Onset |  |  |
| Corticospinal Tract | -0.56 | 0.58 |
| Precentral Gyrus | -1.11 | 0.27 |
| Tapping Rate |  |  |
| Corticospinal Tract (contralateral) | 0.73 | 0.46 |
| Precentral Gyrus | 1.30 | 0.19 |
| ALSFRS-R |  |  |
| Corticospinal Tract | 2.08 | 0.03 |
| Precentral Gyrus | **4.13** | **<0.0001** |
| Frontotemporal Lobe | 0.80 | 0.42 |

### Disease Staging and Projection of Progression in Atrophy

It is not unreasonable to assume that, in ALS, the greatest brain atrophy is found in areas that have been affected the longest by the disease. Using different threshold values on the t-statistics map shown in Figure 1, four stages of the disease were identified based on the brain regions involved and the magnitude of tissue shrinkage (Figure 3). These stages were consistent with the ALS stages defined based on previous pathologic staging studies (Brettschneider *et al.*, 2013). Figure 3 shows the spread of brain atrophy and its correspondence with pathologic stages as defined by Brettschneider et al. (Brettschneider et al. 2013).

**Fig. 3.**
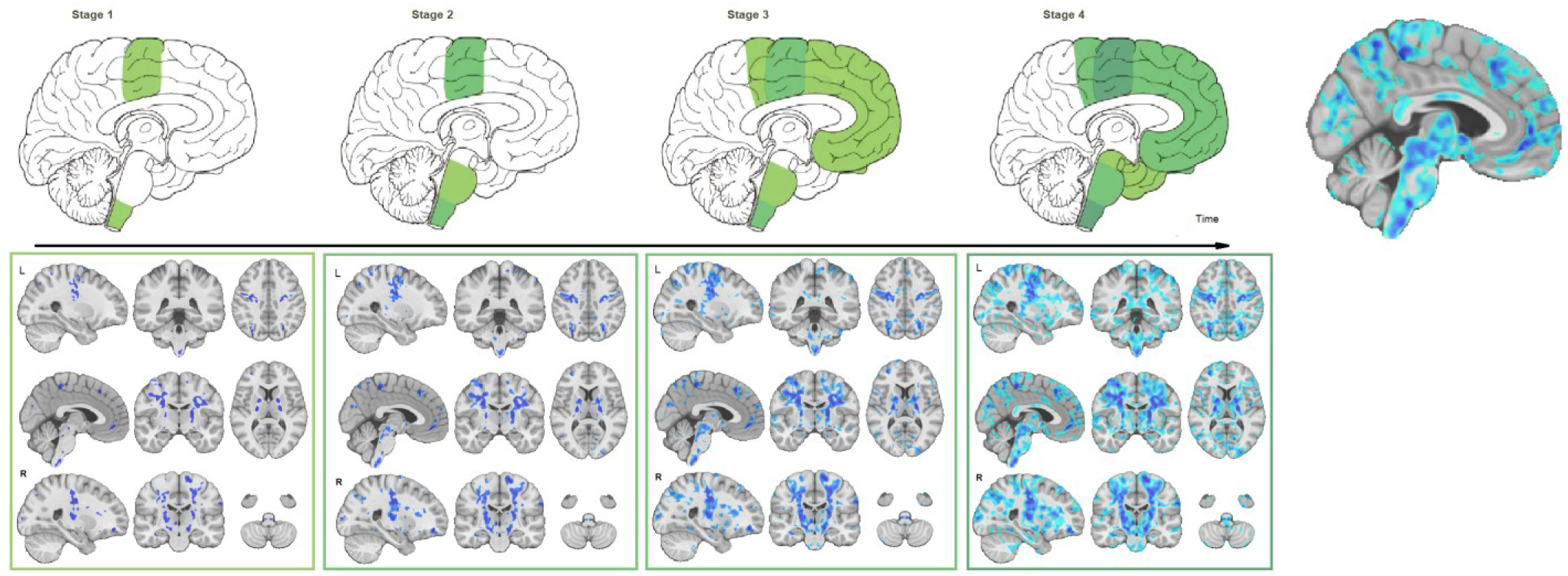
Comparison of disease stages based on magnitude of atrophy and histological stages. The first row indicates the four pathological stages based on Brettschneider et al. (2013) as well as a corresponding sagittal slice from the t-statistics map (Figure 2). The second row shows three sagittal, coronal, and axial slices overlayed with the t-statistics map values thresholded at t-values= −5, −4, −3, and −1.5, to correspond with stages 1-4, respectively.

## Discussion

In this study, using DBM and taking advantage of the large multi-center cohort of ALS patients available from CALSNIC, it was possible to reveal the extensive atrophy pattern characteristic of ALS: bilateral atrophy in the motor cortex and the corticospinal tract including the internal capsule and brainstem, and an overall pattern of ventricular enlargement, along with additional longitudinal atrophy in the bilateral precentral gyri, diffuse white matter, as well as ventricular and cortical sulcus enlargement. The pattern of atrophy described in this paper (Figure 3) is consistent with pathologically defined stages in ALS (Brettschneider et al. 2013). Furthermore, atrophy in the precentral gyrus significantly correlated with disability as assessed with the ALSFRS-R (p<0.0001), similar to a previous study (Sheng *et al.*, 2015).

Previous imaging studies have reported similar patterns of atrophy, although not as comprehensively in general. For example, VBM studies have reported GM reduction in motor, premotor, basal ganglia and frontotemporal regions (Chang *et al.*, 2005; Agosta *et al.*, 2007; Thivard *et al.*, 2007; Agosta *et al.*, 2009; Senda *et al.*, 2011; Sheng *et al.*, 2015; Shen *et al.*, 2016; Buhour *et al.*, 2017; Kim *et al.*, 2017, Chen *et al.*, 2018*b*; Menke *et al.*, 2018). Cortical thickness studies have reported cortical thinning in the motor cortex as well as frontal and temporal areas (Roccatagliata *et al.*, 2009; Verstraete *et al.*, 2012, Chen *et al.*, 2018*c*). A recent meta-analysis reported significant WM reduction in the bilateral supplementary motor areas, precentral gyri, left middle cerebral peduncle, and right cerebellum, involving the corticospinal tract, interhemispheric fibres, subcortical arcuate fibres, and the projection fibres to the striatum and cortico-ponto-cerebral tract (Chen *et al.*, 2018*a*).

The atrophy map obtained in this study provides an *in-vivo* confirmation for the sequentially increasing pathology in alignment with Brettschneider model (Brettschneider et al. 2013). This sequential pattern of disease progression has also been reported in diffusion tensor imaging studies (Kassubek *et al.*, 2014; Müller *et al.*, 2016; Schmidt *et al.*, 2016). Further studies are necessary to determine the nature of the underlying biological changes (microstructural damage, macrostructural atrophy, or a combination) that are observed in T1w and DTI data. To our knowledge, no previous morphometric study of ALS has succeeded to show the correspondence between the pattern of gray and white matter atrophy and disease stages defined by TDP-43 spreading (Brettschneider et al. 2013). Using different threshold values on the t-statistics map (reflecting the severity of atrophy specific to ALS), the four stages described by Brettschneider et al. were identified (Figure 3). The possibility to determine disease stage, and pathological propagation, using structural MRI could have potential implications in the clinical setting. Indeed, using already available and routinely used MRI, non motor progression of symptoms (i.e. cognitive impairment) could be anticipated. Further studies are necessary to investigate the sensitivity of using structural MRI at the level of an individual participant.

The present study has several advantages compared with the previous reports. The multi-center CALSNIC cohort included 66 ALS patients, a larger sample size compared with previous studies (Sheng *et al.*, 2015). The MRI scans were also acquired with higher magnetic field strength (i.e. 3T rather than 1.5T), yielding images with better contrast. All the image processing tools used in this study have been developed and extensively validated for use in multi-center studies involving different MRI systems, and have been used in numerous such studies (Zeighami *et al.*, 2015, Dadar *et al.*, 2018*b*; Misquitta *et al.*, 2018; Dadar *et al.*, 2019; Manera *et al.*, 2019; Sanford *et al.*, 2019). In addition, all analyses included *center* as a categorical random effect to ensure that any residual variability across centers would not bias the results.

We acknowledge that there were limitations to the present study. Not all participants completed all three visits, although the proportion of the number of follow-up visits to baseline were similar across patient and control groups. In addition, the follow-up period (8 months) is relatively short. Future studies investigating DBM changes in ALS with longer follow-ups are warranted, though there are inherent constraints in recruiting such a cohort.

In conclusion, DBM reveals the atrophy pattern characteristic of ALS consistent with previous pathological findings, both in gray and white matter areas and can be used to obtain aquantitative measure of disease burden. DBM measurements might therefore have the potential to be a biomarker in ALS.

## Data availability statement

Anonymized data will be shared at the request of qualified investigators.

## References

Abrahams S, Newton J, Niven E, Foley J, Bak TH. Screening for cognition and behaviour changes in ALS. Amyotroph Lateral Scler Front Degener 2014; 15: 9–14.

Agosta F, Gorno-Tempini ML, Pagani E, Sala S, Caputo D, Perini M, et al. Longitudinal assessment of grey matter contraction in amyotrophic lateral sclerosis: a tensor based morphometry study. Amyotroph Lateral Scler 2009; 10: 168–174.

Agosta F, Pagani E, Rocca MA, Caputo D, Perini M, Salvi F, et al. Voxel-based morphometry study of brain volumetry and diffusivity in amyotrophic lateral sclerosis patients with mild disability. Hum Brain Mapp 2007; 28: 1430–1438.

Andreadou E, Sgouropoulos P, Varelas P, Gouliamos A, Papageorgiou C. Subcortical frontal lesions on MRI in patients with motor neurone disease. Neuroradiology 1998; 40: 298–302.

Avants BB, Epstein CL, Grossman M, Gee JC. Symmetric diffeomorphic image registration with cross-correlation: evaluating automated labeling of elderly and neurodegenerative brain. Med Image Anal 2008; 12: 26–41.

Brettschneider J, Del Tredici K, Toledo JB, Robinson JL, Irwin DJ, Grossman M, et al. Stages of pTDP-43 pathology in amyotrophic lateral sclerosis. Ann Neurol 2013; 74: 20–38.

Brooks BR, Miller RG, Swash M, Munsat TL. El Escorial revisited: revised criteria for the diagnosis of amyotrophic lateral sclerosis. Amyotroph Lateral Scler Other Motor Neuron Disord 2000; 1: 293–299.

Brown J, Deng J, Neuhaus J, Sible IJ, Sias AC, Lee SE, et al. Patient-Tailored, Connectivity-Based Forecasts of Spreading Brain Atrophy. 2019

Brownell B, Oppenheimer DR, Hughes JT. The central nervous system in motor neurone disease. J Neurol Neurosurg Psychiatry 1970; 33: 338–357.

Buhour M-S, Doidy F, Mondou A, Pélerin A, Carluer L, Eustache F, et al. Voxel-based mapping of grey matter volume and glucose metabolism profiles in amyotrophic lateral sclerosis. EJNMMI Res 2017; 7: 21.

Chang JL, Lomen-Hoerth C, Murphy J, Henry RG, Kramer JH, Miller BL, et al. A voxel-based morphometry study of patterns of brain atrophy in ALS and ALS/FTLD. Neurology 2005; 65: 75–80.

Charil A, Corbo M, Filippi M, Kesavadas C, Agosta F, Munerati E, et al. Structural and metabolic changes in the brain of patients with upper motor neuron disorders: a multiparametric MRI study. Amyotroph Lateral Scler 2009; 10: 269–279.

Chen G, Zhou B, Zhu H, Kuang W, Bi F, Ai H, et al. White matter volume loss in amyotrophic lateral sclerosis: A meta-analysis of voxel-based morphometry studies. Prog Neuropsychopharmacol Biol Psychiatry 2018; 83: 110–117.

Chen Z, Liu M, Ma L. Gray Matter Volume Changes over the Whole Brain in the Bulbar- and Spinal-onset Amyotrophic Lateral Sclerosis: a Voxel-based Morphometry Study. Chin Med Sci J 2018; 33: 20–28.

Chen Z, Liu M, Ma L. Cortical Thinning Pattern of Bulbar- and Spinal-onset Amyotrophic Lateral Sclerosis: a Surface-based Morphometry Study. Chin Med Sci J 2018; 33: 100–106.

Coupe P, Yger P, Prima S, Hellier P, Kervrann C, Barillot C. An Optimized Blockwise Nonlocal Means Denoising Filter for 3-D Magnetic Resonance Images. IEEE Trans Med Imaging 2008; 27: 425–441.

Dadar M, Fonov VS, Collins DL, Initiative ADN. A comparison of publicly available linear MRI stereotaxic registration techniques. NeuroImage 2018; 174: 191–200.

Dadar M, Maranzano J, Ducharme S, Collins DL. White matter in different regions evolves differently during progression to dementia. Neurobiol Aging 2019; 76: 71–79.

Dadar M, Zeighami Y, Yau Y, Fereshtehnejad S-M, Maranzano J, Postuma RB, et al. White matter hyperintensities are linked to future cognitive decline in de novo Parkinson’s disease patients. NeuroImage Clin 2018; 20: 892–900.

Fonov V, Evans A, McKinstry R, Almli C, Collins D. Unbiased nonlinear average age-appropriate brain templates from birth to adulthood. NeuroImage 2009; 47: S102.

Gordon PH. Amyotrophic lateral sclerosis: an update for 2013 clinical features, pathophysiology, management and therapeutic trials. Aging Dis 2013; 4: 295.

Hardiman O, Al-Chalabi A, Chio A, Corr EM, Logroscino G, Robberecht W, et al. Amyotrophic lateral sclerosis. Nat Rev Dis Primer 2017; 3: 17071.

Hecht MJ, Fellner F, Fellner C, Hilz MJ, Heuss D, Neundörfer B. MRI-FLAIR images of the head show corticospinal tract alterations in ALS patients more frequently than T2-, T1-and proton-density-weighted images. J Neurol Sci 2001; 186: 37–44.

Kassubek J, Müller H-P, Del Tredici K, Brettschneider J, Pinkhardt EH, Lule D, et al. Diffusion tensor imaging analysis of sequential spreading of disease in amyotrophic lateral sclerosis confirms patterns of TDP-43 pathology. Brain 2014; 137: 1733–1740.

Kato S, Hayashi H, Yagishita A. Involvement of the frontotemporal lobe and limbic system in amyotrophic lateral sclerosis: as assessed by serial computed tomography and magnetic resonance imaging. J Neurol Sci 1993; 116: 52–58.

Kim H-J, Leon M de, Wang X, Kim HY, Lee Y-J, Kim Y-H, et al. Relationship between Clinical Parameters and Brain Structure in Sporadic Amyotrophic Lateral Sclerosis Patients According to Onset Type: A Voxel-Based Morphometric Study. PLOS ONE 2017; 12: e0168424.

Kushner PD, Stephenson DT, Wright S. Reactive astrogliosis is widespread in the subcortical white matter of amyotrophic lateral sclerosis brain. J Neuropathol Exp Neurol 1991; 50: 263–277.

La Joie R, Perrotin A, Barré L, Hommet C, Mézenge F, Ibazizene M, et al. Region-specific hierarchy between atrophy, hypometabolism, and β-amyloid (Aβ) load in Alzheimer’s disease dementia. J Neurosci 2012; 32: 16265–16273.

Landau SM, Mintun MA, Joshi AD, Koeppe RA, Petersen RC, Aisen PS, et al. Amyloid deposition, hypometabolism, and longitudinal cognitive decline. Ann Neurol 2012; 72: 578–586.

Manera AL, Dadar M, Collins DL, Ducharme S, Initiative FLDN. Deformation based morphometry study of longitudinal MRI changes in behavioral variant frontotemporal dementia. NeuroImage Clin 2019: 102079.

Martin JE, Swash M. The pathology of motor neuron disease. In: Motor neuron disease. Springer; 1995. p. 93–118

Menke RAL, Proudfoot M, Talbot K, Turner MR. The two-year progression of structural and functional cerebral MRI in amyotrophic lateral sclerosis. NeuroImage Clin 2018; 17: 953–961.

Misquitta K, Dadar M, Tarazi A, Hussain MW, Alatwi MK, Ebraheem A, et al. The relationship between brain atrophy and cognitive-behavioural symptoms in retired Canadian football players with multiple concussions. NeuroImage Clin 2018; 19: 551–558.

Moradi E, Pepe A, Gaser C, Huttunen H, Tohka J, Initiative ADN, et al. Machine learning framework for early MRI-based Alzheimer’s conversion prediction in MCI subjects. Neuroimage 2015; 104: 398–412.

Müller H-P, Turner MR, Grosskreutz J, Abrahams S, Bede P, Govind V, et al. A large-scale multicentre cerebral diffusion tensor imaging study in amyotrophic lateral sclerosis. J Neurol Neurosurg Psychiatry 2016; 87: 570–579.

Oba H, Araki T, Ohtomo K, Monzawa S, Uchiyama G, Koizumi K, et al. Amyotrophic lateral sclerosis: T2 shortening in motor cortex at MR imaging. Radiology 1993; 189: 843–846.

Ringholz GM, Appel SH, Bradshaw M, Cooke NA, Mosnik DM, Schulz PE. Prevalence and patterns of cognitive impairment in sporadic ALS. Neurology 2005; 65: 586–590.

Roccatagliata L, Bonzano L, Mancardi G, Canepa C, Caponnetto C. Detection of motor cortex thinning and corticospinal tract involvement by quantitative MRI in amyotrophic lateral sclerosis. Amyotroph Lateral Scler 2009; 10: 47–52.

Sanford R, Strain J, Dadar M, Maranzano J, Bonnet A, Mayo NE, et al. HIV infection and cerebral small vessel disease are independently associated with brain atrophy and cognitive impairment. Aids 2019; 33: 1197–1205.

Schmidt R, de Reus MA, Scholtens LH, van den Berg LH, van den Heuvel MP. Simulating disease propagation across white matter connectome reveals anatomical substrate for neuropathology staging in amyotrophic lateral sclerosis. Neuroimage 2016; 124: 762–769.

Senda J, Kato S, Kaga T, Ito M, Atsuta N, Nakamura T, et al. Progressive and widespread brain damage in ALS: MRI voxel-based morphometry and diffusion tensor imaging study. Amyotroph Lateral Scler 2011; 12: 59–69.

Shen D, Cui L, Fang J, Cui B, Li D, Tai H. Voxel-Wise Meta-Analysis of Gray Matter Changes in Amyotrophic Lateral Sclerosis [Internet]. Front Aging Neurosci 2016; 8[cited 2019 Oct 25] Available from: https://www.frontiersin.org/articles/10.3389/fnagi.2016.00064/full

Sheng L, Ma H, Zhong J, Shang H, Shi H, Pan P. Motor and extra-motor gray matter atrophy in amyotrophic lateral sclerosis: quantitative meta-analyses of voxel-based morphometry studies. Neurobiol Aging 2015; 36: 3288–3299.

Sled JG, Zijdenbos AP, Evans AC. A nonparametric method for automatic correction of intensity nonuniformity in MRI data. Med Imaging IEEE Trans On 1998; 17: 87–97.

Strong MJ. Progress in clinical neurosciences: the evidence for ALS as a multisystems disorder of limited phenotypic expression. Can J Neurol Sci 2001; 28: 283–298.

Thivard L, Pradat P-F, Lehéricy S, Lacomblez L, Dormont D, Chiras J, et al. Diffusion tensor imaging and voxel based morphometry study in amyotrophic lateral sclerosis: relationships with motor disability. J Neurol Neurosurg Psychiatry 2007; 78: 889–892.

Verstraete E, Veldink JH, Hendrikse J, Schelhaas HJ, van den Heuvel MP, van den Berg LH. Structural MRI reveals cortical thinning in amyotrophic lateral sclerosis. J Neurol Neurosurg Psychiatry 2012; 83: 383–388.

Waragai M. MRI and clinical features in amyotrophic lateral sclerosis. Neuroradiology 1997; 39: 847–851.

Wilson CM, Grace GM, Munoz DG, He BP, Strong MJ. Cognitive impairment in sporadic ALS: a pathologic continuum underlying a multisystem disorder. Neurology 2001; 57: 651–657.

Yeh F-C, Panesar S, Fernandes D, Meola A, Yoshino M, Fernandez-Miranda JC, et al. Population-averaged atlas of the macroscale human structural connectome and its network topology. NeuroImage 2018; 178: 57–68.

Zeighami Y, Fereshtehnejad S-M, Dadar M, Collins DL, Postuma RB, Mišić B, et al. A clinical-anatomical signature of Parkinson’s Disease identified with partial least squares and magnetic resonance imaging. NeuroImage 2017

Zeighami Y, Ulla M, Iturria-Medina Y, Dadar M, Zhang Y, Larcher KM-H, et al. Network structure of brain atrophy in de novo Parkinson’s disease. eLife 2015; 4: e08440.

